# Monitoring mutant myocilin secretion and localization in trabecular meshwork cell cultures using a protein complementation-based luminescence assay

**DOI:** 10.1101/2025.02.28.640780

**Authors:** Hannah A. Youngblood, Ethan F. Harris, Kaylee P. Lankford, Victoria Garfinkel, John D. Hulleman, Raquel L. Lieberman

## Abstract

Approximately 2-4% of adult onset and 10% of juvenile onset cases of primary open angle glaucoma can be attributed to non-synonymous coding mutations in *MYOC*. One of the key characteristics of a pathogenic *MYOC* mutant is the inability of the resulting protein to be secreted from trabecular meshwork cells. Instead, pathogenic myocilin variants accumulate in the endoplasmic reticulum. Typically, localization of MYOC mutants is compared to wild-type myocilin in cellular secretion assays that use immunoblot to detect myocilin in extracellular media, alongside intracellular soluble and insoluble (aggregated) fractions. Here, we implement a new method that utilizes a complement-based luminescence method in which an 11-residue HiBiT tag is appended to myocilin and complements a truncated nanoluciferase. The method allows for highly sensitive luminescence detection and does not require immunoblot. We tested non-synonymous coding variants T377R, D384G, D395ins, C433Y, T455K, and L486F, in an established immortalized trabecular meshwork cell line. Secretion was tested in 96-well plate format, revealing poor secretion for these mutants compared to wild-type myocilin. For assays conducted in 6-well plates, myocilin mutants were accumulated in intracellular fractions. HiBiT luminescence signals correlated well with immunofluorescence as well as immunoblot but is more sensitive than the latter. Overall, our study demonstrates that complement-based detection of mutant myocilin using luminescence allows for facile and sensitive detection of myocilin localization and has confirmed secretion defects for seven variants.

**Highlights:**

- Mutations in myocilin are causal for early onset open angle glaucoma
- In the lab pathogenic myocilin mutations are characterized by secretion defects
- We validate a complement luminescence assay for detection of myocilin localization
- The luminescence-based assay is more sensitive than traditional immunoblot
- We confirm secretion defects for several mutants not previously characterized

**TOC Graphic:** 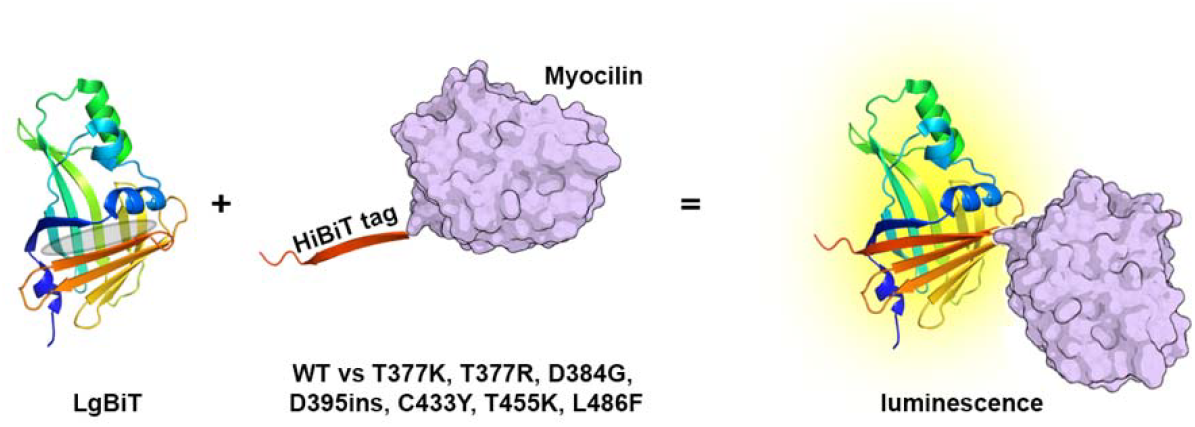

---

Glaucoma is the leading cause of irreversible blindness worldwide with primary open angle glaucoma (POAG) being the most common subtype, affecting ∼60 million people around the globe (Allison et al., 2020). The primary risk factor for glaucoma is elevated intraocular pressure (IOP) (Kwon et al.). IOP elevation is thought to be caused by an imbalance between the production of aqueous humor (AH) from the ciliary body and its outflow through the trabecular meshwork (TM) (Goel et al., 2010). Previous studies have shown that 2-4% of adult-onset POAG cases and 10% of juvenile open angle glaucoma (JOAG) cases are attributed to mutations within myocilin (MYOC), a protein that is highly expressed in the TM. The expression of disease-causing myocilin variants hastens TM dysfunction through a toxic gain-of-function mechanism. Namely, pathogenic variants cannot be secreted from the cell and instead accumulate within the endoplasmic reticulum (ER) causing cytotoxicity (Joe et al., 2003; Kwon et al.; Liu and Vollrath, 2004; Scelsi et al., 2023).

The traditional method for detecting mutant MYOC in a cellular context involves immunoblotting. Cells are transiently transfected with plasmids containing MYOC variants and cultured for approximately 2 days, after which the spent media is collected, and the cells are fractionated into soluble and aggregated fractions (Zhou and Vollrath, 1999). Aggregated MYOC is insoluble in Triton X-100 detergent while MYOC within the detergent-soluble fraction is not aggregated but is otherwise prevented from being secreted. The amount of MYOC contained in each of these 3 fractions is then detected by immunoblot using either an antibody against MYOC or an encoded epitope tag. Detection of intracellular myocilin is important because wild-type myocilin is a secreted protein, and disease-causing mutants exhibit low levels of secretion from cells cultured at 37 °C (Burdon et al., 2022; Gobeil et al., 2004; Jacobson et al., 2001).

In this study, we describe a luciferase complementation reporter assay (NanoBiT (Dixon et al., 2016), **Fig. 1A**) in which a small 11-amino acid HiBit tag is appended to myocilin and complements a truncated nanoluciferase. This small tag allows for sensitive and quantitative detection of MYOC variants in both secreted media and intracellular fractions. The method also overcomes limitations of fusing MYOC to an enhanced Gaussia luciferase (eGLuc2) (Nakahara and Hulleman, 2022; Zadoo et al., 2016). While eGLuc2 allows for quick luciferase-based detection of the reporter protein, it is a large appendage (∼18 kDa), with unknown consequences to MYOC folding.

**Fig. 1.**
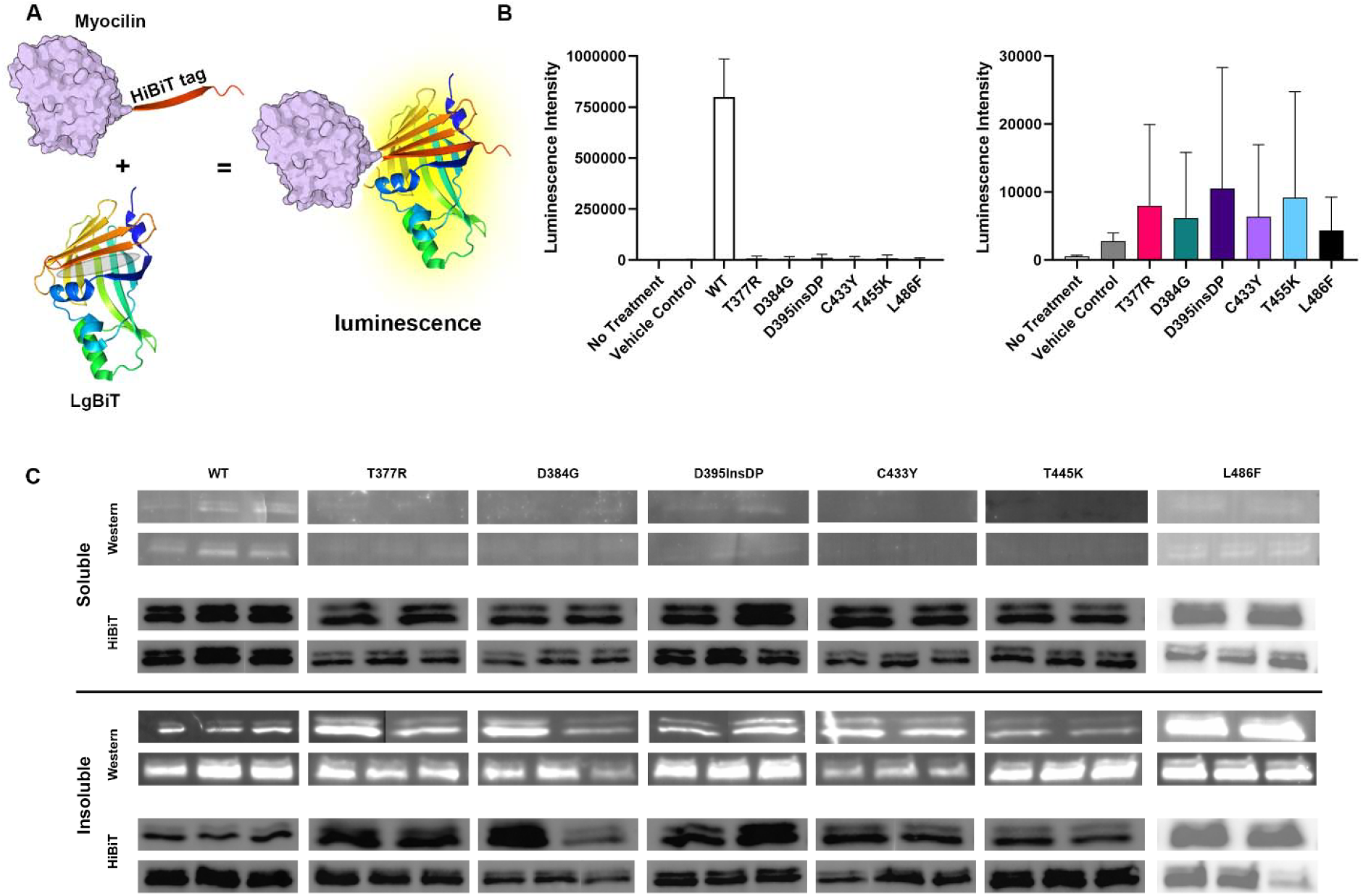
Protein complementation luminescence assay principle and results. (A) Schematic representation of luminescence assay. An 11-amino acid HiBiT tag is appended to myocilin. After culturing cells (not shown), the HiBiT tag completes the missing strand of LgBiT (grey oval) to yield full-length NanoBiT, which produces luminescence upon addition of furimazine. (B) Luminescence intensity of WT and mutant myocilin in secreted media as measured by NanoBiT luminescence assay. Left, the 6 mutants have significantly (p < 0.0001) lower luminescent intensity than WT, suggesting that only WT myocilin is secreted to any extent. Right, rescaling of graph on left with WT removed to allow for comparison of low level luminescence of mutants. Mutants are not significantly different from one another (p > 0.05). (C) Comparison of Western (white bands) and HiBiT blotting (black bands) for soluble and insoluble intracellular fractions of myocilin-expressing TM-1 cells. Technical duplicates or triplicates of two separate transfections are shown for each myocilin variant. Each analysis (Western or HiBiT blot) is conducted on the same respective sample. Overall, a more robust signal was detected for HiBiT compared to Western blot.

Furthermore, the eGLuc2 method was unable to detect intracellular MYOC. As a result, eGLuc2 fusion could not distinguish whether a MYOC mutant found intracellularly was soluble or aggregated.

Using the NanoBiT system, we examined wild-type (WT) myocilin, previously analyzed D395insDP (Nakahara and Hulleman) as a pathogenic positive control variant, and 5 uncharacterized MYOC variants (T377R, D384G, C433Y, T455K, and L486F). T377R (Waryah et al., 2013) involves the same codon as the well-documented and characterized, pathogenic T377M variant (Scelsi et al., 2021). T377R has been detected in multiple generations of a single family with glaucoma, and the disease segregates with the mutation. Variant D384G was found in 5 members of a Chinese Uygar family with POAG (Cai et al., 2012). D395insDP, one of the few documented indel variants of myocilin, was first identified in a Brazilian family with JOAG (Braghini et al., 2013), and then in a case-control study of Brazilian JOAG patients (Svidnicki et al., 2018). C433Y is likewise a rare variant identified via exome sequencing of ∼7000 clinic patients in China, of whom 455 were diagnosed with POAG and the balance had other ocular disorders (Li et al., 2021). T455K (Tian et al., 2007) and L486F (Liao et al., 2016) were each found in large Chinese families; L486F was also found in a large cohort of glaucoma patients in China (Huang et al., 2014; Li et al., 2021) as well as independently in a Finnish family (Liuska et al., 2023).

Plasmids of WT MYOC and MYOC mutants were generated via cloning into the pcDNA 3 vector with C-terminal FLAG (DYKDDDDK) and HiBiT (VSGWRLFKKIS) tags separated by a short valine serine linker (VS) (Dixon et al., 2016; Los et al., 2008; Schwinn et al., 2018). Plasmid fidelity was confirmed by nanopore sequencing (Plasmidsaurus). Plasmids were expressed in immortalized human trabecular meshwork (TM-1) cells, a kind gift from Donna Peters (U. Wisc) (Filla et al., 2002). Cells were grown and maintained in low-glucose (1g/L) Dulbecco’s modified Eagle medium (DMEM, Gibco) supplemented with 10% fetal bovine serum (FBS, Hyclone) and 1-2% penicillin-streptomycin (PS, Gibco) at 37 °C per consensus recommendations (Keller et al., 2018).

Cells were plated at 80-100% confluency 18-24 hours before transfection. For intracellular analysis, experiments were conducted in 6-well plate format whereas for secreted fractions, 96-well plates were used. Transfections (n=2) with respective plasmid DNA were performed using Lipofectamine 2000 (Invitrogen) and 2 μg/well or 100 ng/well for 6-well or 96-well plates. Cells were allowed to incubate for 24 hours before changing the media to serum-free low-glucose DMEM with 1-2% PS. After an additional 24 hours, the media was collected as the secreted MYOC fraction and treated with 1:100 protease inhibitor cocktail (containing cOmplete protease inhibitor (Roche) and phosphatase inhibitor cocktails 2 and 3 (Sigma-Aldrich)). The concentration of total protein in the secreted media was quantified by a BCA Assay (Pierce) according to the manufacturer’s protocol.

The complement luminescence assay was performed on the media (secreted) fraction according to the manufacturer’s recommendations. The LgBiT solution containing a 1:100 dilution of LgBiT protein and a 1:50 dilution of Nano-Glo® HiBiT Lytic Substrate (Promega) was prepared in PBS (Gibco) from the Nano-Glo® HiBiT Lytic Detection System kit (Promega, Cat no. N3030). In a white-walled 96-well plate (ThermoFisher Scientific, Cat no. 165306), 1:1 LgBiT solution was added to the samples. The plate was wrapped in foil and gently rocked for 10 minutes at room temperature. Luminescence readings were obtained using a BioTek Synergy H4 (Agilent) instrument (luminescence filter 460/40 nm). Data analysis was performed using GraphPad Prism software. Statistical significance was tested using one-way ANOVA.

Analysis of the HiBiT luminescence data from the conditioned media shows that all 6 variants secrete to a limited extent but are statistically lower than that of WT (p < 0.0001, **Fig. 1B**). The luminescence between mutants was not statistically different (**Fig. 1B**). Thus, all mutants tested in this experiment exhibit similar levels of compromised secretion in the HiBiT luminescence assay.

Next, we tested whether the soluble and aggregated fractions could be detected with the HiBiT blotting system (Promega, Cat no. N2410) and yield results comparable to those seen with the immunoblot technique. After spent media was collected, adherent cells were lysed using 100 mM Tris-HCl pH 7.4, 3 mM EGTA, 5 mM MgCl_2_, 0.6Mm PMSF, 0.5% Triton-X 100 with 1:100 protease inhibitor cocktail. Lysed cells were scraped, collected, and centrifuged at 10,000 *x g* for 5 minutes to separate the soluble from the aggregated fractions. After the soluble fraction was removed from the insoluble fraction, insoluble samples were washed three-times with ice-cold phosphate-buffered saline (PBS, Gibco) and resuspended in 300 μL of 2x Laemmli buffer with 10% BME. Insoluble samples were then sonicated with a rod sonicator (Qsonica Q125) for 5 minutes with a 10 s on/off pulse at 50% amplitude prior to electrophoresis. The concentration of total protein in the soluble fraction was quantified by a BCA Assay (Pierce) according to the manufacturer’s protocol. Soluble samples were prepared with 2x Laemmli’s to a final concentration of 1x containing 10% β-mercaptoethanol (BME, v/v). All samples containing Laemmli’s buffer were boiled for 10 minutes at 100-110 °C prior to electrophoresis.

HiBiT blotting was performed on the soluble and insoluble fractions according to the manufacturer’s recommendations. Equal micrograms of protein (soluble fraction) or equal volumes (aggregated fraction) were run on polyacrylamide gels in technical duplicate or triplicate; due to the number of samples, multiple gels were required and not all samples were run synchronously. The gels were transferred onto nitrocellulose membranes (BioRad) using a Trans-Blot^®^ Turbo™ Transfer System (BioRad). The membranes were rocked overnight at 4 °C in 1:200 LgBiT protein (Promega, from Cat No. N3030) and a 10x dilution of the Nano-Glo® blotting buffer in 0.1% PBST. The following day, membranes were brought to room temperature and a 1:500 dilution of the Nano-Glo® Luciferase Assay Substrate (Promega, from kit Cat No. N3030) was directly added to the 0.1% PBST solution containing LgBiT. The solution was incubated for 5 minutes. The membrane was then immediately imaged on a ChemiDoc MP Imaging System (BioRad) using the auto-gain chemiluminescence setting.

Similarly, traditional immunoblotting was performed on the soluble and insoluble samples. For immunoblotting, detection using the tag adjacent to HiBiT, namely, FLAG, was used due to their comparable epitope placement in the construct. PVDF membranes (BioRad) were used instead of nitrocellulose. Following transfer, membranes were blocked with 5% milk, washed with 0.1% PBST, incubated with 1:2000 primary antibody (mouse monoclonal Anti-FLAG M2 antibody, Sigma Aldrich AB_259529) for 1 hour, washed with 0.1% PBST, incubated with 1:2000 secondary antibody (goat anti-mouse StarBright B520, Bio-Rad AB_2934034), and washed with PBS before being imaged with the ChemiDoc MP Imaging System.

Given that both the FLAG epitope and HiBiT sequence are linearized in the blotting condition, we initially expected that there would be no significant difference in intensity in the HiBiT versus antibody. However, compared to immunoblot at the same protein concentration, where MYOC was detected only to a low degree in the soluble fractions, NanoBiT luminescence was visible for all samples (**Fig. 1C**). Similarly, insoluble MYOC detection exhibited a similar pattern; even in the cases of low detection, the luminescence readout was more robust than its immunoblotting counterpart. Compared to the soluble samples, the signal for the insoluble samples was more variable, likely because the latter samples are difficult to load onto the gel.

Regardless, our results show that the luminescence detection method yields verifiable signal with significantly higher sensitivity than seen with immunoblotting. The origin of this difference likely lies in the high level of sensitivity of direct luminescence measurements compared to the more indirect antibody-based probe.

As a final confirmation of the fidelity of the HiBiT system to report on myocilin localization, we performed an orthogonal experiment, immunocytochemistry imaging of MYOC in TM-1 cells, using an N-terminal-directed anti-MYOC antibody (Patterson-Orazem et al., 2018). Cells were seeded on glass coverslips (12 mm diameter, Fisher) coated with 2% gelatin (catalog number G1393-20ML, Sigma-Aldrich, St. Louis, MO, USA). The next day, media was changed to serum-free media prior to transfection. Per coverslip, 500 ng of plasmid was complexed with 1.25 µl Lipofectamine 2000 in serum-free media according to the manufacturer’s protocol. Three separate transfections were performed. After 48 hours, cells were washed with 1X PBS (Gibco), fixed with 10% formalin (Fisher Healthcare), permeabilized with 0.03% Triton X-100 (VWR Health Sciences), and blocked in 5% milk before incubating with primary antibodies >12 hours (1:1000 rabbit anti-myocilin, Abcam Ab41552; 1:1000 mouse anti-calnexin, Invitrogen MA3-027). Following a PBS wash, cells were incubated with 1:1000 secondary antibodies (goat anti-rabbit IgG, Alexa Fluor 488, Invitrogen A11034; goat anti-mouse IgG, Cyanine5, Invitrogen A10524) for 1 hour. Following another PBS wash, cells were incubated with 1:1000 Hoechst 33342 (catalog number ENZ51035K100, Enzo Life Sciences, Farmingdale, NY, USA) for ∼30 minutes. Cells were washed before fixing coverslips with ProLongTM Diamond Antifade Mountant (Invitrogen) and allowed to cure for at least 24 hours. Controls included cells treated with media only (no treatment), cells treated with Lipofectamine 2000 only (vehicle control), and cells transfected with αGFP. In addition, negative controls for each condition were treated with only the secondary antibodies and not the primary antibodies to eliminate non-specific antibody detection. Two positively and two negatively stained coverslips were prepared for each condition and 3-4 representative images were acquired per coverslip at 40X magnification using a Leica DMB6 fluorescent microscope (Leica Microsystems, Wetzler, Germany). Brightness and contrast of all images in the panel was equivalently adjusted with Leica Application Suite X software and Microsoft PowerPoint.

Quantification was conducted on unadjusted images using ImageJ by measuring the mean gray value for both MYOC and Hoescht 33342 fluorescence after auto-threshholding with the Triangle auto-threshholding method. The MYOC fluorescence values were normalized to the Hoescht 33342 fluorescence values. Normalized values were averaged per replicate and condition. Average values were background-subtracted with the average value for the no treatment control. Fold change was calculated relative to normalized, background-subtracted WT fluorescence. Data were analyzed by one-way ANOVA (GraphPad Prism) of the average background-subtracted, normalized intensity of all representative images per sample.

Despite a relatively low transfection efficiency, as shown by use of a green fluorescence protein (GFP) transfection control plasmid, transfection yielded noticeable expression and immunocytochemistry-based fluorescence of all plasmids (**Fig. 2A-B**). As expected, WT myocilin was expressed diffusely throughout the cell with occasional perinuclear banding (**Fig. 2B)**. By contrast, but in line with luminescence and immunoblot data above (**Fig. 1B-C**), all MYOC mutants were predominantly localized within dense perinuclear bands (**Fig. 2B**). Furthermore, the number of cells per field of view with intracellular myocilin was noticeably higher for the mutants versus WT myocilin. This difference is reflected by a higher fluorescence intensity for all MYOC mutants as compared to WT, although this difference was not statistically significant (**Fig. 2C**). Furthermore, the prevalence of a dense perinuclear band in cells transfected with MYOC mutants as opposed to the more sparse and diffuse expression of WT MYOC suggests that the mutants may be aggregating, an observation consistent with HiBiT blot and Western blot results (**Fig. 1C**). Of the mutants, L486F had the fluorescence intensity closest to WT, which corroborates our immunoblot and HiBiT blot results (**Fig. 1C, 2C**). The reason for this finding is unclear but we speculate that L486F may accumulate to a lower extent than other mutants and instead may be partially degraded. By contrast, D384G exhibited the highest fluorescence intensity in immunocytochemistry, whereas the strongest signal in Western and HiBiT blots were for D395insDP (**Fig. 1C, 2C**). This latter discrepancy can be explained by the fact that fluorescence intensity of immunocytochemistry data measures all intracellular MYOC, which includes both the soluble and aggregated fractions. Related, the aggregated species in D395insDP may be less accessible to antibody detection in the context of immunocytochemistry compared to denaturing conditions of a blot.

**Fig. 2.**
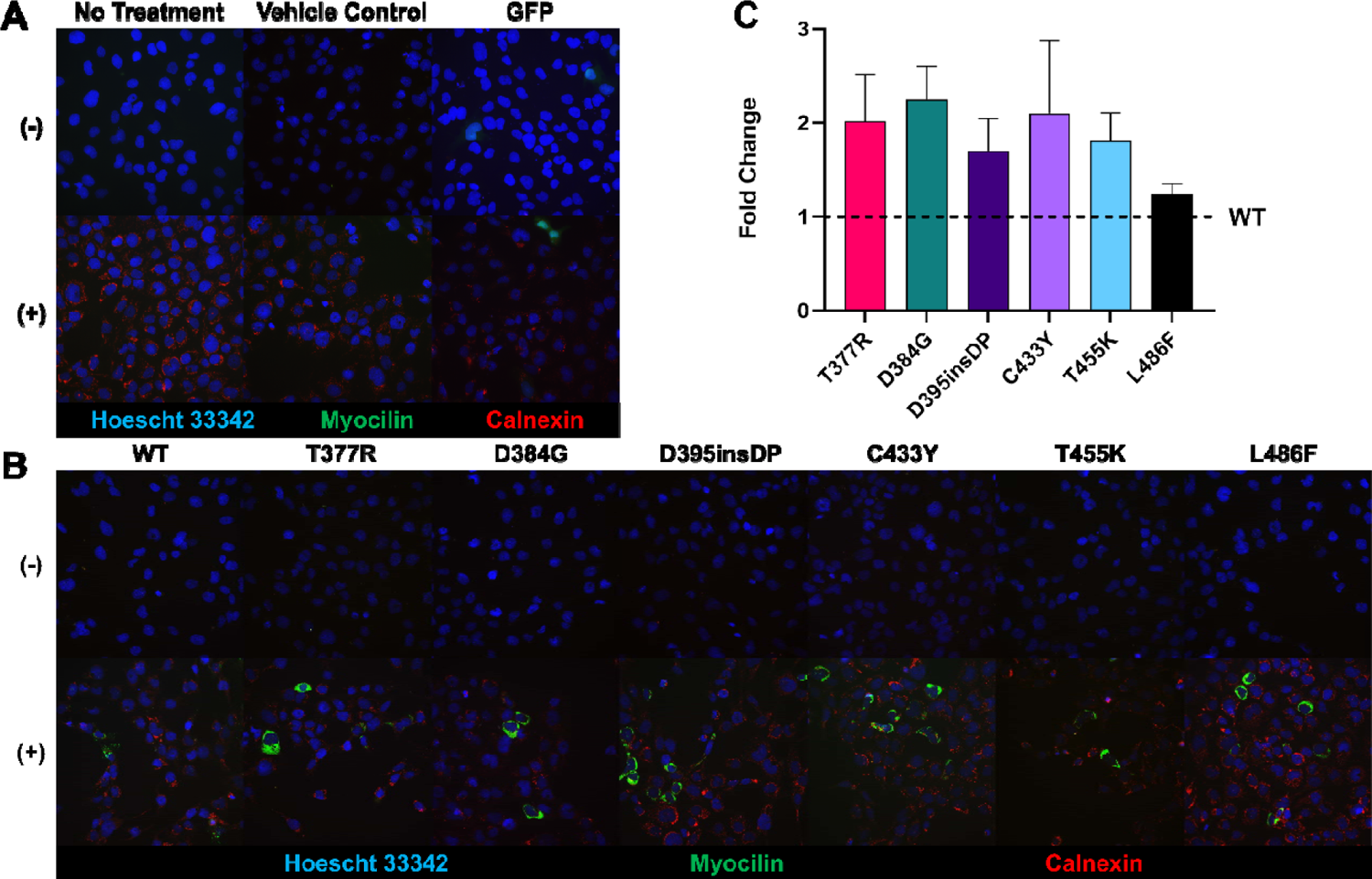
Immunofluorescence imaging of HiBiT-tagged myocilin variants expressed in TM-1 cells. (A) Immunocytochemistry of no treatment, vehicle, and GFP controls. GFP serves as a transfection control, showing relatively low transfection. Calnexin serves as a marker for the ER. (-) indicates negative stain controls (secondary antibody only), and (+) indicates cells stained with both primary and secondary antibodies. Images are representative of 3 independent transfections. (B) Immunocytochemistry of WT myocilin and 6 myocilin mutants. WT myocilin exhibits low fluorescence likely because it is secreted (see Fig. 1B). (C) Quantification of immunofluorescent imaging. Signal intensity is higher for all mutants compared to wild-type, in line with intracellular sequestration detected in Fig. 1C.

In summary, we have shown that the complement-based luminescence assay that appends an 11-residue HiBiT tag to the C-terminus of MYOC can readily detect mutant MYOC in both extracellular and intracellular fractions (**Fig. 1A**). Further, we have confirmed secretion defects for six misfolding variants of MYOC that, to our knowledge, have not been investigated previously in the laboratory. This assay overcomes the previous limitation of the eGLuc2 fusion whereby only secreted myocilin could be detected and required a large ∼18 kDa reporter protein appendage (Zadoo et al., 2016). Detection of the intracellular fraction is both more sensitive and notably less laborious than immunoblot, though it is still a multi-week process that requires the addition of a protein tag and is therefore not readily translated to a clinical setting. It may be possible to further optimize this assay for direct luminescence measurements of the intracellular fractions, since no cross-reactivity with LgBiT was observed in the HiBiT blots. It may also be possible to improve quantification by comparing luminescence to a standard. In terms of limitations of the assay, any future interest in evaluating truncation mutations would require the HiBiT tag to be moved to the N-terminus of myocilin. In the long term, a more streamlined evaluation of how well a given new MYOC variant is secreted can support efforts in the clinic to genotype individuals and assess likelihood of developing glaucoma. This may enable early monitoring and disease intervention, which is critical to slowing disease progression and potentially preventing disability due to vision loss (de Vries et al., 2024; Zebardast and Wiggs, 2024).

## Author contributions

HAY-Formal analysis, investigation, methodology, writing original draft, writing-review and editing

EFH-Investigation, formal analysis, writing-original draft

KLP, VG – MYOC HiBiT variant generation and validation

JDH-Conceptualization, project administration, formal analysis, funding acquisition, methodology, writing-original draft, writing-review and editing

RLL-Project administration, formal analysis, funding acquisition, methodology, writing-original draft, writing-review and editing

